# The broad spectrum mixed-lineage kinase 3 inhibitor URMC-099 prevents acute microgliosis and cognitive decline in a mouse model of perioperative neurocognitive disorders

**DOI:** 10.1101/537472

**Authors:** Patrick Miller-Rhodes, Cuicui Kong, Gurpreet S. Baht, Ramona M. Rodriguiz, William C. Wetsel, Harris A. Gelbard, Niccolò Terrando

**Author notes:** shared correspondence.

## Abstract

Perioperative neurocognitive disorders (PND), including delirium and postoperative cognitive dysfunction, are serious complications that afflict up to 50% of surgical patients and for which there are no disease-modifying therapeutic options. Here, we test whether prophylactic treatment with the broad spectrum mixed-lineage kinase 3 inhibitor URMC-099 prevents surgery-induced neuroinflammation and cognitive impairment in a translational model orthopedic surgery-induced PND. We used a combination of two-photon scanning laser microscopy and CLARITY with light-sheet microscopy to define surgery-induced changes in microglial morphology and dynamics. Orthopedic surgery induced microglial activation in the hippocampus and cortex that accompanied impairments in episodic memory. URMC-099 prophylaxis prevented these neuropathological sequelae without impacting bone fracture healing. Together, these findings provide compelling evidence for the advancement of URMC-099 as a therapeutic option for PND.

## Background

Delirium and postoperative cognitive dysfunction, now collectively referred to as perioperative neurocognitive disorders (PND), have become the most common complications after routine surgical procedures (1, 2). Almost half of surgical patients experience cognitive disturbances following surgery, which can lead to poorer prognosis and a higher 1-year mortality rate in subjects with pre-existing neurodegeneration (3). Currently, there are no approved disease-modifying therapeutic options to treat PNDs and the pathogenesis of these complex phenomena is not fully understood.

To clarify the pathophysiology of PNDs, we developed an orthopedic surgery mouse model that recapitulates clinical orthopedic procedures that associate with neurological disorders in frail subjects (4). In this model, we and others have described a key role for neuroinflammation, including changes in microglial and astroglial morphology, and hippocampal-dependent memory impairments (5–8). Orthopedic surgery-induced cognitive impairment depends on the presence of microglia, as their depletion abrogates these deficits in mice (9). In support of these findings, a recent study identified a correlation between microglial activation and cognitive impairment in patients receiving intra-abdominal surgery using positron emission tomography imaging (10).

We have demonstrated the therapeutic efficacy of the small-molecule, blood-brain barrier (BBB)-permeable, mixed-lineage kinase 3 (MLK3) inhibitor URMC-099 in preclinical models of Alzheimer’s disease, HIV-1-associated neurocognitive disorders, and multiple sclerosis (11–13). Despite its development as a MLK3 inhibitor (14), URMC-099 also inhibits multiple kinase targets that mediate pathological signaling cascades in innate immune cells and end-organ target cells (such as neurons) (13, 15). However, while URMC-099 can reduce untoward microglial responses in multiple neuroinflammatory settings, its effects on fracture healing and bone remodeling during perioperative recovery are unknown. Here, we test the hypothesis that prophylactic URMC-099 treatment can attenuate neuroinflammation and cognitive impairment without interfering with fracture healing.

## Materials & Methods

### Animals

Male C57BL/6 and CX3CR1-GFP mice were obtained from Jackson Laboratories (Bar Harbor, ME). Heterozygous male and female CX3CR1-GFP mice (Jax stock no. 005582), at 3 months of age, were used for the longitudinal 2PLSM experiment. Male C57BL/6 mice (Jax stock no. 000664), at 3 and 9 months of age, were used for the remainder of the experiments All animal procedures were carried out under protocols approved by Duke University Medical Center and University of Rochester Medical Center, Institutional Animal Care and Use Committee under the National Research Council Guide for the Care and Use of Laboratory Animals, 8^th^ edition. Both Duke University and University of Rochester Medical Center are AAALAC accredited institutions.

### Orthopedic surgery model

Orthopedic surgery was performed as described previously (4). Briefly, the left hind limb was subjected to an open tibial fracture with intramedullary fixation under general anesthesia (isoflurane: 4% induction, 2% maintenance) and analgesia (buprenorphine: 0.1 mg/kg, s.c.). Sham-treated mice were administered anesthesia and analgesia as above but without surgical intervention.

### URMC-099 administration

URMC-099 (M.W. 421) was synthesized as originally described in Goodfellow et al. (14). URMC-099 drug solutions were prepared by dissolving 20 mg of URMC-099 in 0.5 mL sterile dimethyl sulfoxide (DMSO; D8779, Sigma-Aldrich, St. Louis, MO). We then added 4 mL polyethylene glycol 400 (PEG400; 91893-250-F, Sigma-Aldrich) and 5.5 mL sterile saline (National Drug Code NDC0409-4888-10). The final concentration was 2 mg/mL URMC-099 in a 10 mL volume. The vehicle was the same solution minus URMC-099. For the experiments described in Figure 1, mice were administered (i.p.) 3 injections at 12 hour intervals prior to orthopedic surgery and two injections post-surgery at identical intervals. For all other experiments, mice were treated only with 3 injections (spaced 12 hr apart) prior to surgery. Drug solutions were coded such that experimenters were blinded to the experimental conditions for the duration of the experiments.

**Figure 1.**
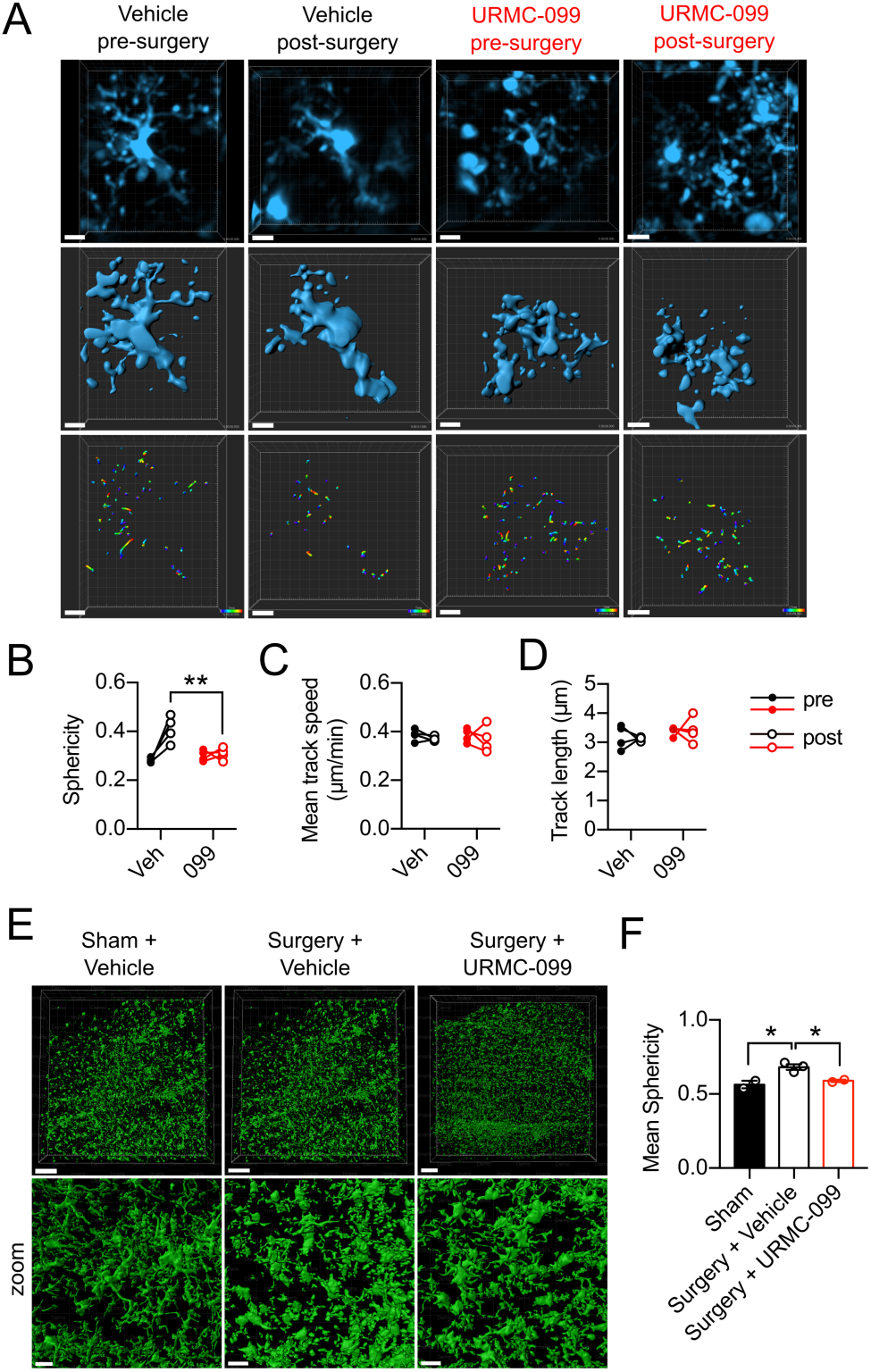
URMC-099 prophylaxis *prevents* microglial morphological changes following orthopedic surgery. 3-month old mice received three doses i.p. of URMC-099 (10 mg/kg) prior to undergoing sham or orthopedic surgery. **A**, Representative cropped and drift-corrected XYZ stacks obtained by 2PLSM (top panels), Imaris 3D surface reconstructions (middle panels), and process movement tracks (bottom panels) (scale bar = 10 μm). **B**, Orthopedic surgery increased microglial sphericity in vehicle-treated, but not URMC-099-treated, mice. No differences were observed for mean track speed, **C**, nor mean track length, **D**. **E**, Representative 3D surface reconstructions of light-sheet micrographs obtained from optically cleared hippocampal sections (scale bar = 500 μm, top panels; scale bar = 100 μm, bottom panels). **F**, Mean microglial sphericity was increased by orthopedic surgery and URMC099 treatment prevented this. N=4 (B-D), N=2-3 (F); results presented as mean ± SEM; *P<0.05, **P<0.01, versus surgery + vehicle group; repeated-measures ANOVA (B-D) or one-way ANOVA (F) with Dunnett’s multiple comparison test.

### Piezoelectric thinned-skull cortical window surgery

The thinned skull cortical window (TSCW) procedure was performed using a piezoelectric surgical technique (17). Piezoelectric surgery was performed using a piezosurgery handpiece (Mectron) fitted with an OT11 osteotomy insert (Mectron) under constant, manual irrigation (1 mL/min) with ice-cold aCSF to minimize the heating of the bone during the thinning process. A cover glass was glued over the thinned area of the skull and a custom head-plate was placed with dental cement to facilitate repeated intravital imaging. Isoflurane was used for anesthesia (4% induction, 2% maintenance) during the TCSW procedure. The TCSW surgery was limited to 30 min or less to minimize the mouse’s exposure to variables that might confound this study’s results (e.g., duration under anesthesia).

### Two-photon intravital microscopy

A custom two-photon laser-scanning microscope (2P-LSM) was used for intravital imaging (Ti:Sapphire, Mai-Tai, Spectraphysics; modified Fluoview confocal scan head, ×20 lens, 0.95 numerical aperture, Olympus). Two-photon excitation was achieved using 100-fsec laser pulses (80MHz) tuned to 840nm with a power of ∼50mW measured at the sample after the objective lens. For all experiments, a 565 nm dichroic mirror was used in conjunction with 580/180 (GFP) and 605/55 (rhodamine B) filters. Rhodamine B dextran (2% w/v, 75 μl total) was injected retroorbitally after ketamine (100 mg/kg)/xylazine (10mg/kg) anesthesia and 5 minutes prior to intravital imaging. Mice were maintained at 37°C during intravital imaging and recovery. Importantly, intravital imaging sessions were limited to 40 minutes per mouse to avoid re-exposure to ketamine/xylazine anesthesia. Intravital imaging was performed using 4X digital zoom and at a 512 × 512 pixel resolution. Stacks containing 41 slices (1μm step size) were imaged every 90 seconds for 15 minutes to obtain the XYZT stacks used for analysis.

### CLARITY tissue processing and light-sheet microscopy

Brains were harvested following transcardial perfusion with 30 to 50 mL of PBS (#10010-023; Gibco) and 20 to 30 mL 4% paraformaldehyde (PFA) (#158127; Sigma-Aldrich) in PBS (pH 7.4) under isoflurane anesthesia. After a post-fixation step (24 hr in 4% PFA at 4°C), brains were sectioned by vibratome into 1 mm-thick coronal sections. Sections were incubated in 1x PBS overnight at 4°C prior to polymerization and tissue clearing. Polymerization and tissue clearing were accomplished using the X-CLARITY™ reagents and equipment (Logos Biosystems). Sections were first saturated in 1mL hydrogel solution (X-CLARITY™ Hydrogel Solution Kit, #C1310X, Logos Biosystems) at 4°C for 24h and were subsequently incubated in the polymerization system for 3 hr at 37°C. Polymerized sections were then actively cleared in electrophoretic tissue clearing (ETC) solution (#C13001, Logos Biosystems) using the X-CLARITY™ Tissue Clearing System with the following settings: 0.9 A, 37°C, for 3h. Cleared sections were washed overnight three times with 0.2% Triton-X100 (T8787, Sigma) in PBS on a shaker at room temperature. Immunohistochemistry was performed by incubating cleared tissue sections with rabbit anti-Iba1 (1:300; cat no. 019-19741; Wako) in 10% normal donkey serum + 0.2% Triton-X100 in PBS (blocking buffer) at 4°C for 3 days. Following three wash steps (0.1% Triton-X100 in PBS, overnight at 4°C), sections were incubated with Alexa Fluor 594-conjugated donkey anti-rabbit secondary antibody (1:500) and DAPI diluted in blocking buffer for 3 days at 4°C. Sections were subsequently stored in PBS until mounting.

For imaging, sections were mounted in 1% agarose (#BP160-100, FisherScientific) and hung in 1mL syringes (#300013, BD Biosciences) without tips followed by incubation with mounting solution (#C13101, Logos Biosystems) at 4°C overnight. Samples were placed at room temperature for 1 hr before imaging. Imaging was performed using a light sheet fluorescence microscope (LSFM, Z1, Zeiss, Germany) at the Duke University Light Microscopy Core Facility. Imaging was performed by suspending samples with mounting solution in a CLARITY-optimized sample chamber and orienting the samples such that they were parallel to the detection lens. XYZ stacks were acquired at 1920 × 1920 pixel resolution and 16-bit depth using 5x or 20x objectives. The refractive index of the detection objective was set as 1.46. A 405/488/561/640 filter set was used for laser excitations with laser intensities set between 0.5 and 2%. An exposure time of 29.97 ms for each frame was used and a step size of 0.659 μm was used. Three-dimensional surface reconstructions of microglia were achieved using Imaris as described below.

### Imaris Image Analysis

Individual microglia were cropped from XYZT stacks, registered, and drift-corrected using ImageJ. Individual microglia XYZT stacks were imported into Imaris (Bitplane) for analysis. For motility analysis, the “Spots” function was used to assign spots to individual microglial processes. Spots were assigned automatically using an estimated diameter of 2 µm, which were subsequently filtered by “Quality” and again manually to remove Spots that did not correspond to the individual microglia under analysis. The “Connected Components” algorithm was then used to generate the movement tracks that were quantified. Movement tracks with a duration of less than 3min were excluded from analysis. For morphological analysis, the “Surface” function was used. Surface detail was set to 0.691 microns. Surfaces were then generated by manual thresholding and were manually filtered to exclude surfaces that did not correspond to the individual microglia under analyses. Sphericity — defined as the ratio of the area of a sphere (with the same volume as the given particle) to the surface area of the particle — was used to quantify microglial morphology. Similar measures have been used previously to quantify changes in microglia morphology, albeit in 2-dimensions (28). Three-dimensional surface reconstructions of microglia were also performed on XYZ stacks derived from CLARITY tissue sections using the “Surface” function as described above with the following modifications: individual microglia were not cropped (instead, whole XYZ stacks were reconstructed) and surface detail was set to 2 µm to facilitate image processing.

### Behavioral Testing

Mice used for behavioral experiments were all housed in a temperature and humidity controlled environment, in Optimice ventilated cages (Animal Care Systems, Centennial, CO) with free access to water and mouse chow (5R58 Mouse Pico Diet, LabDiet, St. Louis, MO) under a 14 hr light: 10 hr dark light cycle with the onset of light at 0700 hrs. For all behavioral tests, mice were transferred to the test room immediately following surgery and were housed in this room for the duration of the study. All test rooms were maintained under the same conditions as the colony housing room. Two memory tasks were administered to assess different aspects of memory performance 24 hours following orthopedic surgery. The first task, a “What-Where-When” Object Recognition Test (21) was used to evaluate different aspects of episodic memory. This task involves three phases designated “Set A”, “Set B”, and “Challenge.” Mice were habituated to a 60 × 40 × 24 cm arena for 5 min. For Set A mice were returned to the arena to explore four identical objects placed in configuration A (Figure 2A) for 5 min. Animals were removed and placed into their home-cages for an inter-trial interval (ITI) of 50-55 min. For Set B mice returned to the arena and were exposed to configuration B objects (Figure 2A) for 5 min. The Challenge trial was administered after a 50-55 min ITI and included two Set A and two B objects arranged in a sequence reflecting the “What”, “Where”, and “When” aspects of episodic memory (Figure 2A). The actual objects used are depicted in Supplementary Figure S2A. All behavioral trials were recorded by a video camera suspended above the test arena that was interfaced to a computer running Media Player II (Noldus Information Technology, Asheville, NC). Videos were scored in a blinded fashion using behavioral recognition and nose-point tracking with Ethovision 11 (Noldus Information Technology, Asheville, NC). These variables included duration of contacts within each object zone (contact = nose within 1 cm of the physical object and with the body axis oriented toward the object). “What”, “Where”, and “When” memory indices were computed from exploration times in the Challenge trial. The “What” aspect of episodic memory was defined as the mean exploration time for both A objects minus the mean exploration time for both B objects over the mean exploration time of all objects. The “What” aspect takes advantage of the animal’s preference for exploring the less recently experienced (A objects) over those more recently experienced (B objects). The “Where” aspect was defined as the mean exploration time for displaced object A minus the mean exploration time for stationary object A over the mean exploration time of both A objects. The “When” aspect was defined as the mean exploration time for “oldest” stationary object A minus the mean exploration time for the two most recent B objects over the mean exploration time for all three of these objects. The “When” index takes advantage of mice exploring the less recent “A” object over the more recent “B” objects while these objects all remain in their original locations.

**Figure 2.**
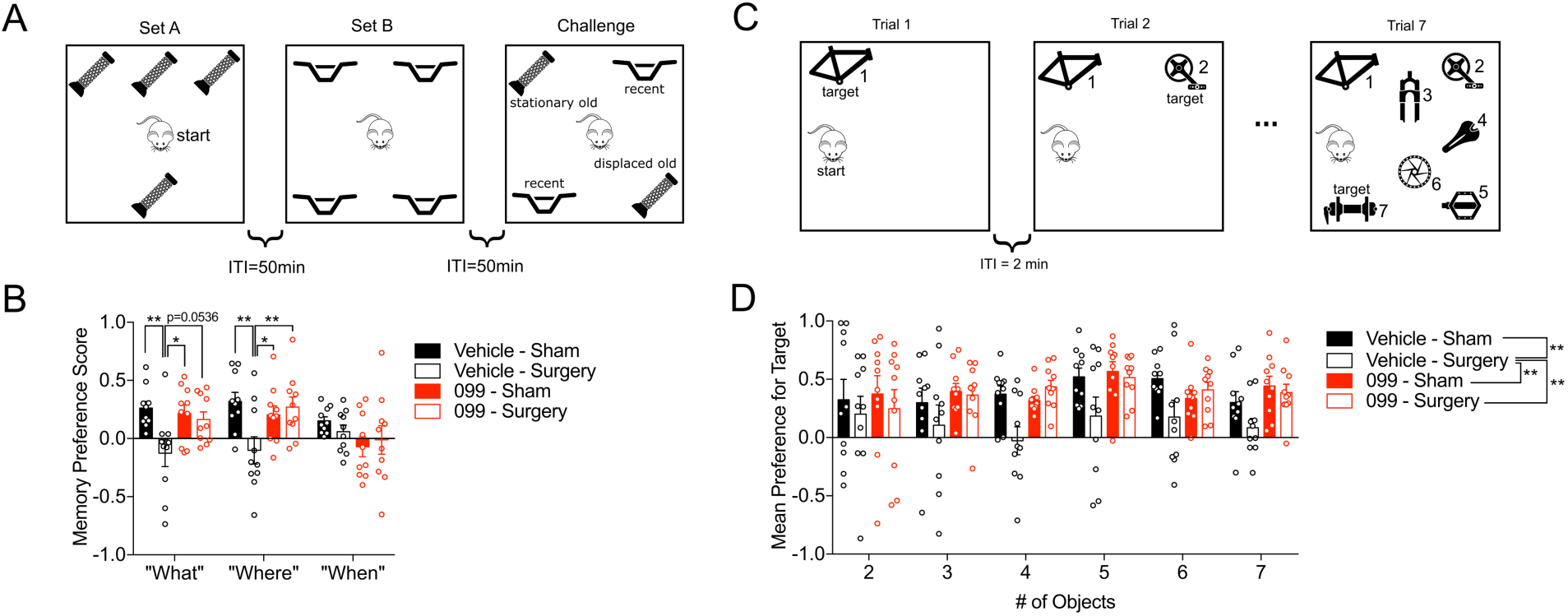
URMC-099 prophylaxis abrogates orthopedic surgery-induced memory deficits. 3-month old mice received three doses i.p. of URMC-099 (10 mg/kg) prior to undergoing sham or orthopedic surgery. **A**, An illustration of the “What-Where-When” Object Discrimination test. **B**, URMC-099 prevented the surgery-induced impairments in the “What” and “Where” phases of the test, whereas “When” was unaffected. **C**, An illustration of the Memory Load Object Discrimination test. **D**, Overall URMC-099 prevented the surgery-induced impairments in the Memory Load test. N=10; results presented as mean ± SEM; *P<0.05, **P<0.01, versus surgery + vehicle group; repeated-measures ANOVA with Dunnett’s multiple comparison test.

Memory load was evaluated using the Memory Load Object Discrimination test (22). This task was conducted in a 42 × 42 × 20 cm arena and it consisted of 7 trials (ranging from 2-5 min, depending on the number of objects present); each trial was separated by an ITI of 1-2 min. On each subsequent trial, a unique novel object was placed into the arena together with the previously presented objects (i.e., the “familiar” objects). Thus, the arena contained one object in trial 1, two objects in trial 2, etc. The objects used in the test are depicted in Supplementary Figure S3A, B. Trial lengths were increased as the number of objects was increased: 2 min (trials 1-3), 3 min (trials 4-5), and 5 min (trials 6-7). For each trial, the newly added object was designated the “target” object. Time spent exploring the target object versus all other objects was recorded as described above for the “What-Where-When” task. Memory preference scores for each trial were computed as ratios of the mean exploration time for the target object minus the mean exploration time for all other objects over the mean exploration time for all objects.

### Analysis of Fracture Healing

Fracture calluses were assessed as previously described (29). Samples were dissected 21 days after injury and fixed in 10% Zn-formalin at room temperature for 5 days. Micro-computed tomography (microCT) analysis was conducted using a Scanco vivaCT 80 (Scanco Medical, Brüttisellen, Switzerland) at a scan resolution of 8µm. Calluses were scanned 1mm proximal and 1mm distal from the fracture site and assessed for total volume (TV) and bone volume (BV) in mm^3^, and bone mineral density (BMD) in mg HA/mm^3^. Fixed fracture calluses were decalcified using 12% EDTA pH 7.4, cleared of EDTA, and embedded into paraffin. Sections were cut at a thickness of 5µm and stained using safranin-O/fast-green to visualize bone and cartilage. A minimum of five sections were used to conduct computer-assisted histomorphometry analysis and results were presented as an amount relative to the total area of the fracture callus.

### Statistics

All data were analyzed by Graphpad PRISM (GraphPad Software, San Diego, CA) and represented as mean ± SEM. Bone healing data were analyzed by unpaired, two-tailed t-tests. The remainder of the data were analyzed by one-way or repeated-measures ANOVA with Dunnett’s multiple comparison test as indicated in the text for each experiment. Statistical significance is defined as P < 0.05 throughout.

## Results & Discussion

Microgliosis and BBB opening accompany cognitive deficits in mice after orthopedic surgery (4–7, 16). We evaluated whether URMC-099 treatment could reverse these pathological changes in a cohort of 9-month old male mice. Mice were treated with three intraperitoneal (i.p.) injections of URMC-099 (10 mg/kg) prior to and two injections over the next day following surgery, with all injections spaced 12 hours apart. At 24 hours post-surgery, hippocampal microgliosis was observed in vehicle-treated mice receiving orthopedic surgery relative to sham-treated controls using stereological analyses of F4/80+ cells (P = 0.0003; Supplementary Figure S1A, B). In contrast, URMC-099-treated mice exhibited a significant reduction (P = 0.0003) in the density of F4/80+ cells in the hippocampus relative to the sham-treated controls (Supplementary Figure S1B), indicating that our treatment paradigm effectively inhibited surgery-induced hippocampal microgliosis. BBB integrity was examined also using IgG immunostaining. We observed significantly less IgG leakage in URMC-099-treated mice receiving surgery and sham-treated controls as compared to vehicle-treated mice receiving surgery (p-values ≤ 0.044; Supplementary Figure S1C). Hence, URMC-099 was efficacious in preventing both microgliosis and leakage through the BBB.

We next tested whether URMC-099 prophylaxis — pre-treatment with three injections of URMC-099 (10 mg/kg, i.p.), spaced 12 hr apart, with the last dose occurring an hour before surgery — is sufficient to *prevent* microglial activation following orthopedic surgery. We used longitudinal two-photon laser-scanning microscope (2P-LSM) to define surgery-induced changes in microglial physiology and to test URMC-099’s ability to prevent these changes. Longitudinal 2P-LSM was performed by imaging vehicle or URMC-099-treated mice immediately prior to and 24 hours post-orthopedic surgery and was achieved using a modified thin-skull cortical window (TSCW) technique (17). Images from vehicle-treated mice exhibited a reduction in the process complexity of microglia post-surgery compared to their pre-surgery images, as quantified by cell “sphericity” (Figure 1A, B). Notably, URMC-099 prophylaxis abrogated this post-surgical effect, as microglia from URMC-099-treated mice remained mostly unchanged between their pre- and post-surgery images (Figure 1A, B). This result was replicated on a more global scale using light-sheet microscopy of CLARITY-processed hippocampal sections (Figure 1E, F). We evaluated also microglial process motility using our longitudinal 2P-LSM paradigm. However, we detected no significant differences in microglial process motility between pre- and post-surgery images irrespective of treatment (Figure 1C, D). Although others have observed increases in microglial process motility following i.p. lipopolysaccharide (LPS) injection (18), this report has been contradicted by other work demonstrating no changes (19) or decreased motility (20) under similar, albeit not identical, conditions. Alternatively, because genetic knockout of *Tlr4* does not prevent cognitive decline in mice receiving orthopedic surgery (7), it is possible that surgical trauma and peripheral LPS injection elicit distinct effects on microglial physiology.

We evaluated hippocampal-dependent memory in our orthopedic surgery model using two behavioral tasks: the “What-Where-When” task (21) and the Memory Load Object Discrimination task (22). We tested for cognitive impairment beginning 24 hours post-surgery. As such, our behavioral paradigm models acute, delirium-like cognitive deficits, similar to those observed in a prospective cohort study of elderly orthopedic surgical patients (23). The “What-Where-When” task tests various components of episodic memory in mice by requiring mice to remember which objects they explored (“What”), their object location (“Where”), and the temporal phase when they were explored (“When”) (Figure 2A; Supplementary Figure S2A). Orthopedic surgery induced memory impairments in vehicle-treated mice in the “What” (P = 0.005) and “Where” (P = 0.007) aspects of the task (Figure 2B). Additionally, we found that URMC-099 prevented the surgery-induced memory impairment in the “Where” (P = 0.0183) and “What” (P = 0.0432) aspects of the task (Figure 2B). No differences in performance were detected among our experimental conditions in the “When” aspect of the task, but this could be due to the low levels of performance across all groups in this task. Importantly, since the object contact times were similar across groups at the each of the three phases of the task, the deficiencies of the vehicle sham group on the “What” and “Where” phases of the task cannot be attributed to an inability to interact with objects (Supplementary Figure S2B). Effects of surgery on motor performance in this task were examined also. Surprisingly, both the vehicle- and URMC-099-treated groups that received orthopedic surgery traveled over greater distances than their sham counterparts (P-values≤0.006) during the Set A phase of the task (Supplementary Figure S2C). Collectively, the object contact and motor performance results indicate that orthopedic surgery does not adversely affect the ability of mice to perform the necessary locomotion to interact with objects in this task.

We further evaluated memory processes using a novel Memory Load Object Discrimination task (22) (Supplementary Figure S3). In this task, mice are presented with a single object during the first trial, and then additional objects are subsequently added to the testing arena on each subsequent trial until the mouse encounters a total of seven objects (Figure 2C). After trial 1, each subsequent trial tests the mouse’s ability to discriminate a novel object from the familiar object(s). Hence, this test evaluates the how many items the mouse can hold in its memory before it can no longer recognize the novel object.

Overall, vehicle-treated mice that received surgery performed significantly worse overall relative to the vehicle-treated (P = 0.003) and URMC-099-treated (p =0.0015) sham controls (Figure 2D). Importantly, URMC-099 treatment prevented memory load impairment in mice receiving surgery (P = 0.0025) (Figure 2D). Taken together, our results indicate that surgery-induced impairments in hippocampal-dependent memory can be prevented by pretreatment with URMC-099.

Anti-inflammatory drugs often impair fracture healing, limiting their translational potential for treating PND. This issue arises because fracture healing depends upon pro-inflammatory cytokines (24) and the recruitment of inflammatory cells that mediate fracture repair (25). To evaluate whether prophylactic URMC-099 impairs fracture healing in surgically-treated mice, we analyzed fracture calluses harvested 21 days post-surgery with microCT and histomorphophometry analyses. MicroCT found no differences in bone mineralization between the surgical vehicle and surgical URMC-099 groups (Figure 3A-C) and histomorphometry analysis also confirmed an absence of group differences in bone deposition (Figure 3D-E). URMC-099 directly interferes with pro-inflammatory cytokine expression by inhibiting kinases that mediate this process (14, 15). The lack of effect of URMC-099 on bone healing suggests that it does not completely abrogate inflammatory responses and may permit certain beneficial inflammatory processes to occur. Alternatively, it is possible that anti-inflammatory drugs that *do* interfere with bone healing (e.g., nonsteroidal anti-inflammatory drugs) exert their detrimental effects through multiple mechanisms that extend beyond the initial mobilization of innate immunity, as has been suggested by others (26). Finally, although MLK3 activation is known to regulate bone mineralization (27), we contend that our prophylactic, time-limited, treatment paradigm helped to preclude any detrimental effect of our drug on long-term bone repair.

**Figure 3.**
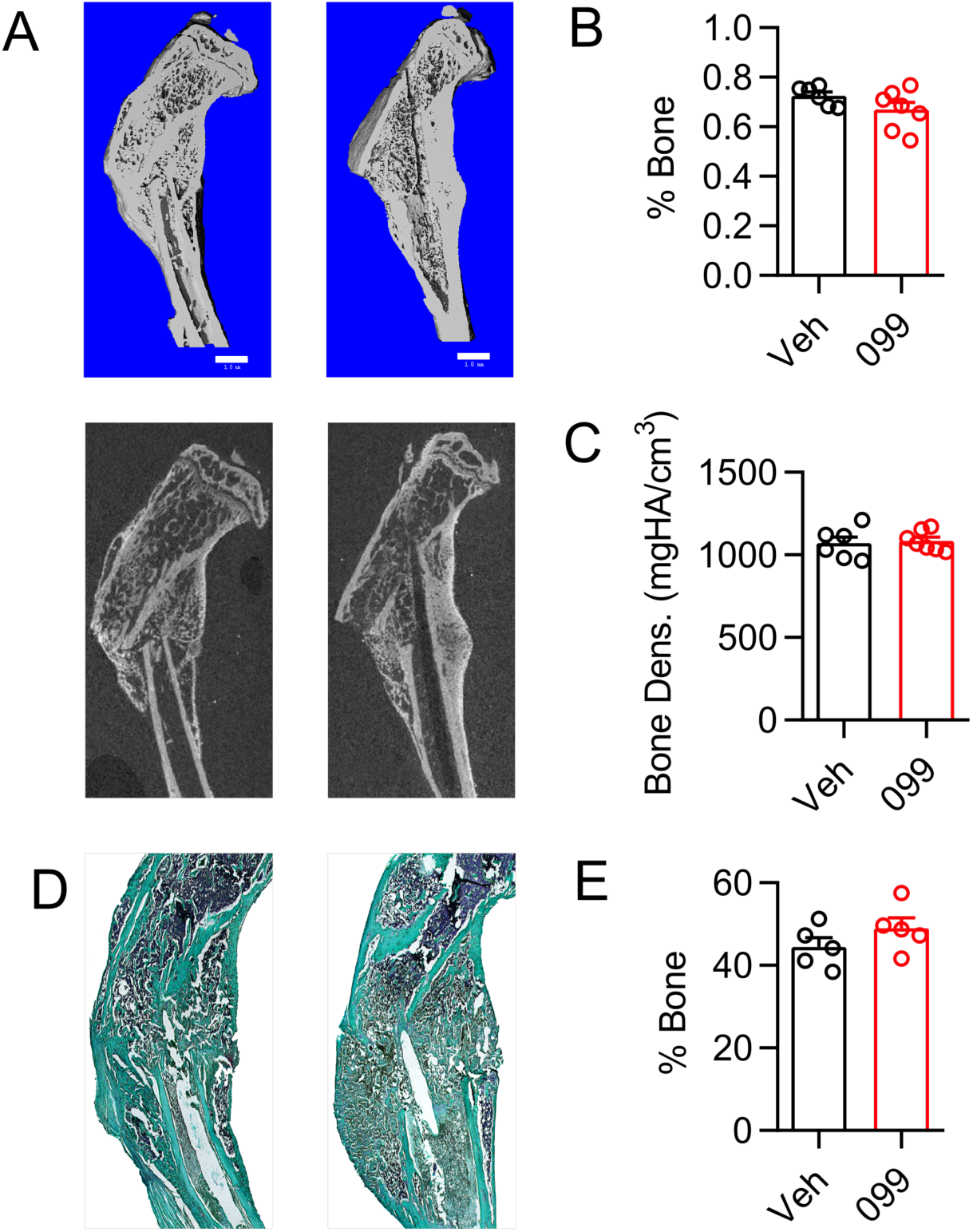
URMC-099 pre-treatment does not alter bone fracture healing. 3-month old male mice were treated with either with URMC-099 or vehicle and fractures were induced surgically. Fracture calluses were investigated 21 days post-fracture to assess bone healing. **A-C**, MicroCT was used to assess bone volume per total callus volume and bone mineral density. **D-E**, Histomorphometry was used to determine the percent bone tissue within the fracture callus. N=6-7; results presented as mean ± SEM. Data analyzed by unpaired two-tailed t-test.

## Conclusions

In conclusion, the present study further defines the neurocognitive sequelae that accompany orthopedic surgery. Because URMC-099 inhibits multiple kinase targets, we cannot ascertain which of its targets are primarily responsible for mediating the therapeutic effects we observed. However, our recently published findings in a mouse model of experimental autoimmune encephalitis indicate that URMC-099’s broad-spectrum activity may be advantageous to the narrow-spectrum profile of a more highly-selective MLK3 inhibitor (13). Thus, inhibiting multiple kinase signaling pathways may be necessary to effect disease-modifying outcomes for neuroinflammatory conditions, including PND.

## Abbreviations

BBB: blood-brain barrier
ip: intraperitoneal
LPS: lipopolysaccharide
MLK3: mixed-lineage kinase 3
PND: perioperative neurocognitive disorders
TSWC: thin-skull cortical window
2P-LSM: two-photon laser-scanning microscope

## Acknowledgments

We wish to thank Mr. Christopher Means of the Duke University Mouse Behavioral and Neuroendocrine Analyses Core Facility for his care of the animals and diligent work during behavioral testing; Dr. Ping Wang in the Terrando lab for helping with the CLARITY samples; and Monique Mendes in Ania Majewska’s lab (URMC) for helping with the acquisition of the 2-photon microscopy data. Design, synthesis and validation of URMC-099 was supported, in part, by NIH grants P01MH64570 and RO1 MH104147 (HAG). Some of the software used in the behavioral testing was purchased with a grant from the North Carolina Biotechnology Center. The study was supported by the NIH RO1 AG057525 (NT).

## Disclosures

URMC-099 is owned by URMC (HA Gelbard. lead inventor: patent nos. US 8,846,909 B2; 8,877,772; and 9,181,247 and associated international patents). HAG has no commercial disclosures regarding URMC-099 at present.

---

**Figure S1.**
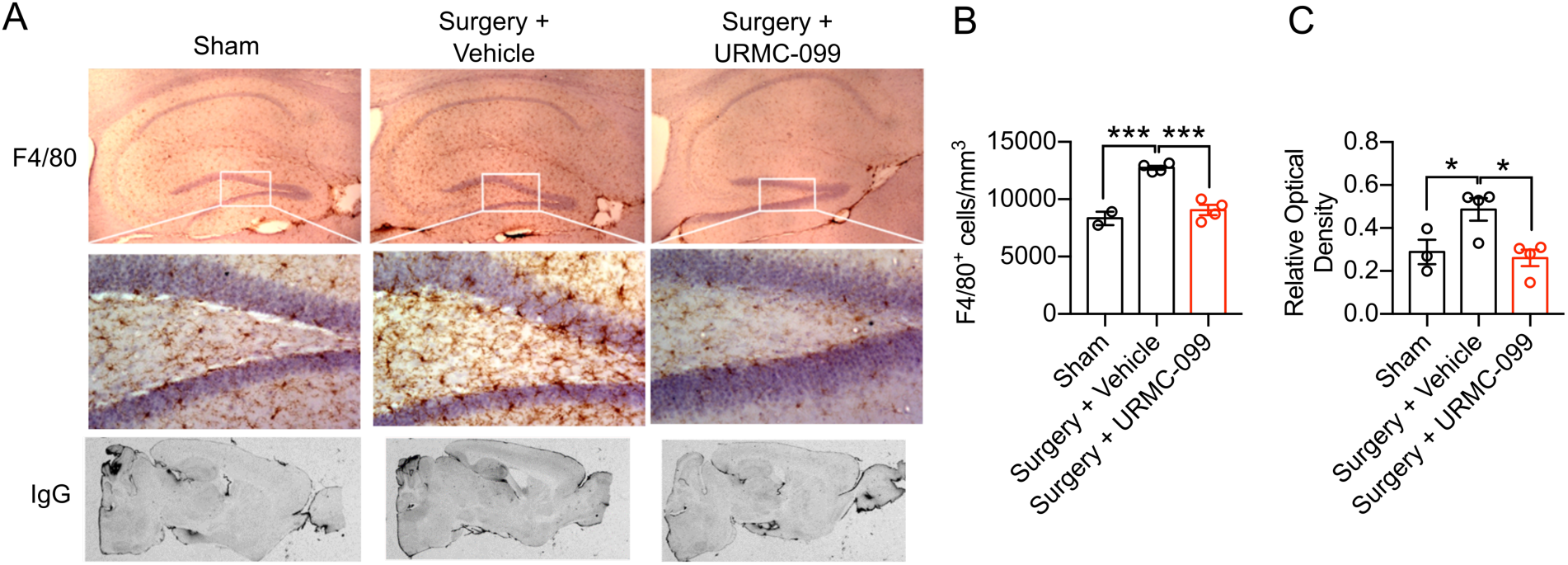
URMC-099 treatment *reverses* microgliosis and blood brain barrier leak following orthopedic surgery. 9-month old male mice underwent open tibial fracture and were terminated 24hr surgery. URMC-099 was given i.p. at 10 mg/kg, 3 doses before surgery and two doses after surgery based on Marker et al. 2013. ***A***, representative images of F4/80 microglial/macrophage staining (top panels) and IgG staining (bottom panels) show that URMC-099 significantly attenuated microgliosis activation in the hippocampus. **B**, Differences in microglia cell numbers were quantified by unbiased stereology. **C**, IgG relative optical density was quantified to measure differences in blood brain barrier opening. N=2-4; results presented as mean ± SEM; *P<0.05, ***P<0.001, versus surgery + vehicle group; one-way ANOVA with Dunnett’s multiple comparison test.

**Figure S2.**
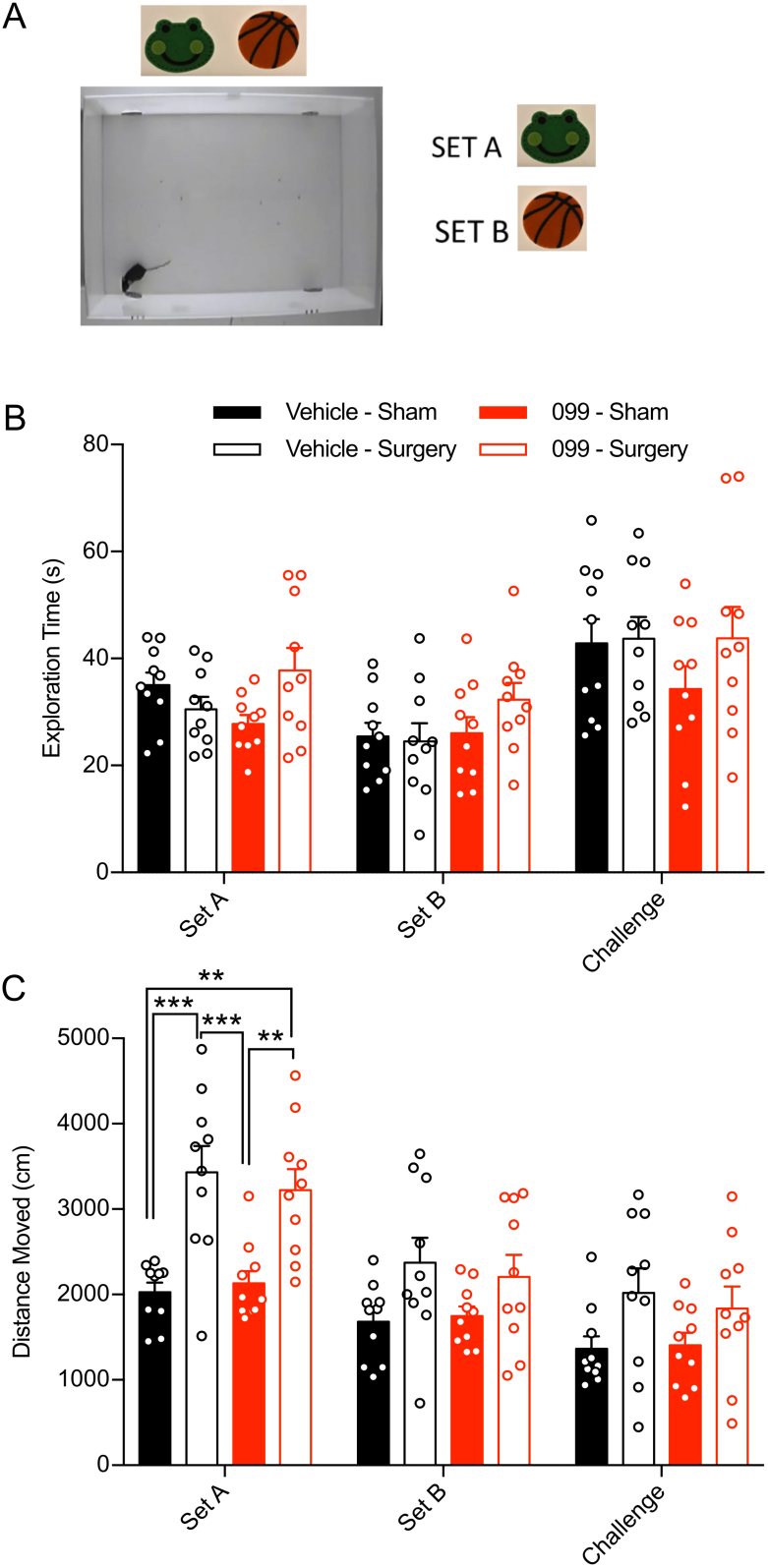
Orthopedic surgery does not impair the mice’s ability to interact with objects but it does affect their locomotion. Three-month old mice received three injections of 10 mg/kg URMC-099 (i.p.) prior to undergoing sham or orthopedic surgery. The “What-Where-When” task was administered 24 hours post-surgery. **A**, Illustration depicting the arena and objects for Set A and Set B used in the task. **B**, Neither surgery nor drug treatment affected the total exploration times throughout the task. **C**, Orthopedic surgery increased the total distance traveled during “Set A” relative to sham-treated controls for both the vehicle- and URMC-099-treated mice. N=10, results presented as means ± SEM; *P<0.05, ***P<0.001, analyzed by repeated-measures ANOVA with Tukey’s multiple comparison test.

**Figure S3.**
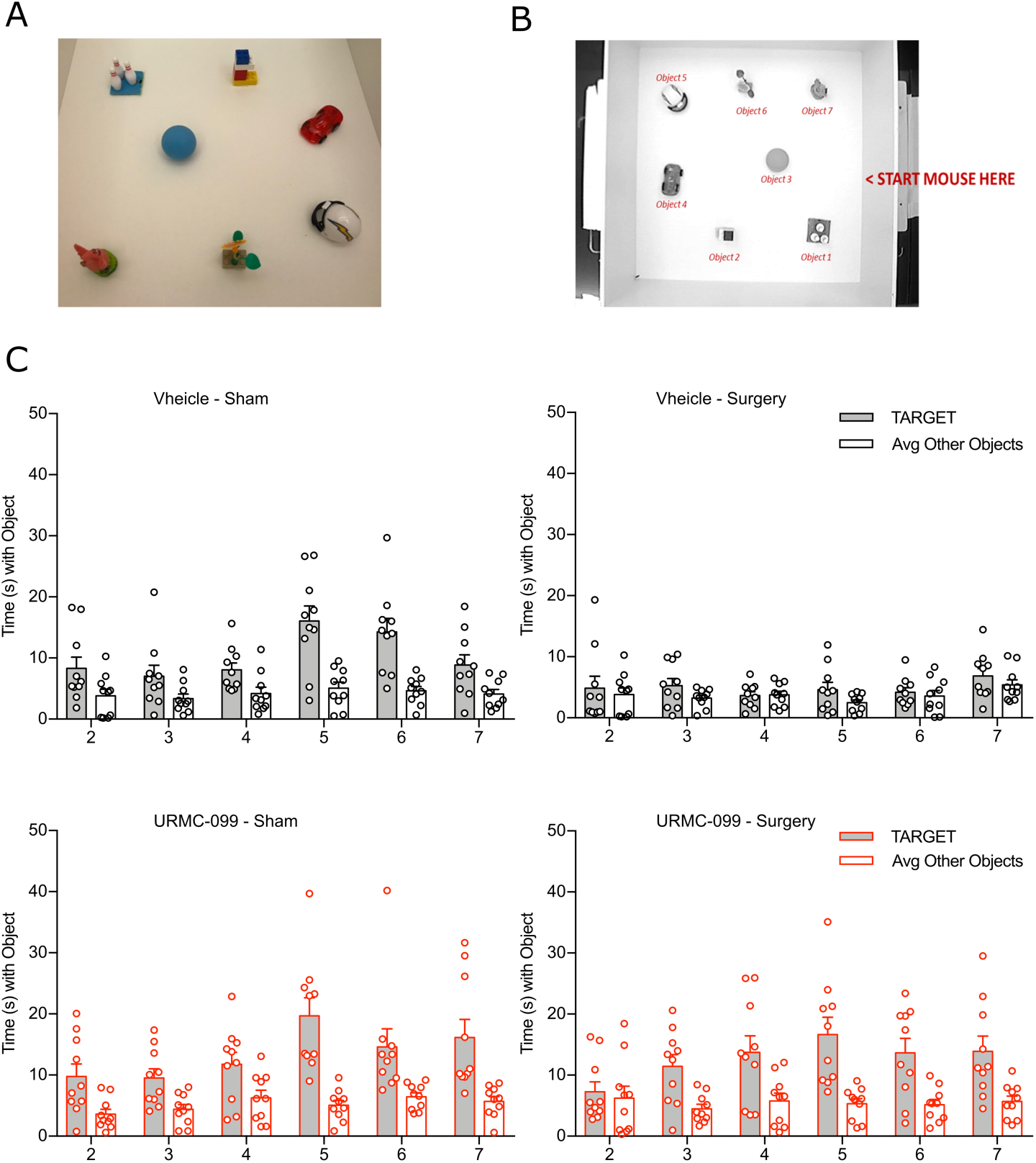
Exploration times for the target versus the other objects in the memory load task. Three-month old mice received three injections of 10 mg/kg URMC-099 (i.p.) prior to undergoing sham or orthopedic surgery. Behavioral tasks were administered beginning 24 hours post-surgery. **A**, Illustration depicting the objects used for the task. **B**, Illustration indicating the object positions with the mouse starting position for the test. **C**, Trial exploration times for each treatment condition. N=10, results presented as means ± SEM.

## References

1. Ags/Nia Delirium Conference Writing Group PC, Faculty. The American Geriatrics Society/National Institute on Aging Bedside-to-Bench Conference: Research Agenda on Delirium in Older Adults. J Am Geriatr Soc. 2015;63(5):843–52.

2. Evered L, Silbert B, Knopman DS, Scott DA, DeKosky ST, Rasmussen LS, et al. Recommendations for the nomenclature of cognitive change associated with anaesthesia and surgery-2018. Br J Anaesth. 2018;121(5):1005–12.

3. Lee HB, Oldham MA, Sieber FE, Oh ES. Impact of Delirium After Hip Fracture Surgery on One-Year Mortality in Patients With or Without Dementia: A Case of Effect Modification. Am J Geriatr Psychiatry. 2017;25(3):308–15.

4. Xiong C, Zhang Z, Baht GS, Terrando N. A Mouse Model of Orthopedic Surgery to Study Postoperative Cognitive Dysfunction and Tissue Regeneration. J Vis Exp. 2018(132).

5. Cibelli M, Fidalgo AR, Terrando N, Ma D, Monaco C, Feldmann M, et al. Role of interleukin-1beta in postoperative cognitive dysfunction. Ann Neurol. 2010;68(3):360–8.

6. Terrando N, Gomez-Galan M, Yang T, Carlstrom M, Gustavsson D, Harding RE, et al. Aspirin-triggered resolvin D1 prevents surgery-induced cognitive decline. FASEB J. 2013;27(9):3564–71.

7. Terrando N, Monaco C, Ma D, Foxwell BM, Feldmann M, Maze M. Tumor necrosis factor-alpha triggers a cytokine cascade yielding postoperative cognitive decline. Proc Natl Acad Sci U S A. 2010;107(47):20518–22.

8. Zhang MD, Barde S, Yang T, Lei B, Eriksson LI, Mathew JP, et al. Orthopedic surgery modulates neuropeptides and BDNF expression at the spinal and hippocampal levels. Proc Natl Acad Sci U S A. 2016;113(43):E6686–E95.

9. Feng X, Valdearcos M, Uchida Y, Lutrin D, Maze M, Koliwad SK. Microglia mediate postoperative hippocampal inflammation and cognitive decline in mice. JCI Insight. 2017;2(7):e91229.

10. Forsberg A, Cervenka S, Jonsson Fagerlund M, Rasmussen LS, Zetterberg H, Erlandsson Harris H, et al. The immune response of the human brain to abdominal surgery. Ann Neurol. 2017;81(4):572–82.

11. Dong W, Embury CM, Lu Y, Whitmire SM, Dyavarshetty B, Gelbard HA, et al. The mixed-lineage kinase 3 inhibitor URMC-099 facilitates microglial amyloid-beta degradation. J Neuroinflammation. 2016;13(1):184.

12. Kiyota T, Machhi J, Lu Y, Dyavarshetty B, Nemati M, Zhang G, et al. URMC-099 facilitates amyloid-beta clearance in a murine model of Alzheimer’s disease. J Neuroinflammation. 2018;15(1):137.

13. Bellizzi MJ, Hammond JW, Li H, Gantz Marker MA, Marker DF, Freeman RS, et al. The Mixed-Lineage Kinase Inhibitor URMC-099 Protects Hippocampal Synapses in Experimental Autoimmune Encephalomyelitis. eNeuro. 2018;5(6).

14. Goodfellow VS, Loweth CJ, Ravula SB, Wiemann T, Nguyen T, Xu Y, et al. Discovery, synthesis, and characterization of an orally bioavailable, brain penetrant inhibitor of mixed lineage kinase 3. J Med Chem. 2013;56(20):8032–48.

15. Marker DF, Tremblay M-È, Puccini JM, Barbieri J, Gantz Marker MA, Loweth CJ, et al. The new small-molecule mixed-lineage kinase 3 inhibitor URMC-099 is neuroprotective and anti-inflammatory in models of human immunodeficiency virus-associated neurocognitive disorders. The Journal of neuroscience : the official journal of the Society for Neuroscience. 2013;33(24):9998–10010.

16. Terrando N, Eriksson LI, Ryu JK, Yang T, Monaco C, Feldmann M, et al. Resolving postoperative neuroinflammation and cognitive decline. Ann Neurol. 2011;70(6):986–95.

17. Clark SD, Mikofsky R, Lawson J, Sulzer D. Piezo High Accuracy Surgical Osteal Removal (PHASOR): A Technique for Improved Cranial Window Surgery in Mice. J Vis Exp. 2018(133).

18. Gyoneva S, Davalos D, Biswas D, Swanger SA, Garnier-Amblard E, Loth F, et al. Systemic inflammation regulates microglial responses to tissue damage in vivo. Glia. 2014;62(8):1345–60.

19. Kondo S, Kohsaka S, Okabe S. Long-term changes of spine dynamics and microglia after transient peripheral immune response triggered by LPS in vivo. Mol Brain. 2011;4:27.

20. Paris I, Savage JC, Escobar L, Abiega O, Gagnon S, Hui CW, et al. ProMoIJ: A new tool for automatic three-dimensional analysis of microglial process motility. Glia. 2018;66(4):828–45.

21. DeVito LM, Eichenbaum H. Distinct contributions of the hippocampus and medial prefrontal cortex to the “what-where-when” components of episodic-like memory in mice. Behav Brain Res. 2010;215(2):318–25.

22. Sannino S, Russo F, Torromino G, Pendolino V, Calabresi P, De Leonibus E. Role of the dorsal hippocampus in object memory load. Learn Mem. 2012;19(5):211–8.

23. Witlox J, Slor CJ, Jansen RW, Kalisvaart KJ, van Stijn MF, Houdijk AP, et al. The neuropsychological sequelae of delirium in elderly patients with hip fracture three months after hospital discharge. Int Psychogeriatr. 2013;25(9):1521–31.

24. Cho TJ, Gerstenfeld LC, Einhorn TA. Differential temporal expression of members of the transforming growth factor beta superfamily during murine fracture healing. J Bone Miner Res. 2002;17(3):513–20.

25. Claes L, Recknagel S, Ignatius A. Fracture healing under healthy and inflammatory conditions. Nat Rev Rheumatol. 2012;8(3):133–43.

26. Ramirez-Garcia-Luna JL, Wong TH, Chan D, Al-Saran Y, Awlia A, Abou-Rjeili M, et al. Defective bone repair in diclofenac treated C57Bl6 mice with and without lipopolysaccharide induced systemic inflammation. J Cell Physiol. 2019;234(3):3078–87.

27. Zou W, Greenblatt MB, Shim JH, Kant S, Zhai B, Lotinun S, et al. MLK3 regulates bone development downstream of the faciogenital dysplasia protein FGD1 in mice. J Clin Invest. 2011;121(11):4383–92.

28. Madry C, Kyrargyri V, Arancibia-Carcamo IL, Jolivet R, Kohsaka S, Bryan RM, et al. Microglial Ramification, Surveillance, and Interleukin-1beta Release Are Regulated by the Two-Pore Domain K(+) Channel THIK-1. Neuron. 2018;97(2):299–312 e6.

29. Baht GS, Nadesan P, Silkstone D, Alman BA. Pharmacologically targeting beta-catenin for NF1 associated deficiencies in fracture repair. Bone. 2017;98:31–6.

